# Silencing of ATP synthase β reduces phytoplasma multiplication in the leafhopper vector *Euscelidius variegatus*

**DOI:** 10.1101/2020.10.02.324350

**Authors:** Luciana Galetto, Simona Abbà, Marika Rossi, Matteo Ripamonti, Sabrina Palmano, Domenico Bosco, Cristina Marzachì

## Abstract

The leafhopper *Euscelidius variegatus* is a natural vector of the chrysanthemum yellows phytoplasma (CYp) and a laboratory vector of the Flavescence dorée phytoplasma (FDp). Previous studies indicated a crucial role for insect ATP synthase α and β subunits during phytoplasma infection of the vector species. Gene silencing of ATP synthase β was obtained by injection of specific dsRNAs in *E. variegatus.* Here we present the systemic and long-lasting nature of such silencing, its effects on the small RNA profile, the significant reduction of the corresponding protein expression, and the impact on phytoplasma acquisition capability. The specific transcript expression was silenced at least up to 37 days post injection with an average reduction of 100 times in insects injected with dsRNAs targeting ATP synthase β (dsATP) compared with those injected with dsRNAs targeting green fluorescent protein (dsGFP), used as negative controls. Insects injected either with dsATP or dsGFP successfully acquired CYp and FDp during feeding on infected plants. However, the average phytoplasma amount in dsATP insects was significantly lower than that measured in dsGFP specimens, indicating a probable reduction of the pathogen multiplication rate when ATP synthase β was silenced. The role of the insect ATP synthase β during phytoplasma infection process is discussed.

## 1. Introduction

Phytoplasmas are plant pathogenic bacteria associated with diseases that cause severe economic impacts on many crops worldwide (Tomkins et al., 2018). These Mollicutes are currently classified within the provisional genus *‘Candidatus* Phytoplasma’ (Marcone, 2014). They are transmitted by phloem feeding hemipteran insects (leafhoppers, planthoppers, and psyllids) in a persistent propagative manner (Bosco and Marzachì, 2016). Phytoplasmas are obligate parasites that live and multiply in close association with their hosts: in plants they inhabit phloem elements (Marcone, 2014), in insects they enter the anterior midgut and colonize salivary glands to be re-transmitted (Koinuma et al., 2020). Control strategies to counteract phytoplasma diseases are mainly based on removal of infected plants and insecticide treatments against vectors, with undesired side effects on environment and public health (Jarausch and Torres, 2014). Moreover, the unculturable nature of these pathogens hampered the studies of their basic biology and their pathogenicity mechanisms. In such a scenario, RNA interference (RNAi) could be exploited as a useful tool for species-specific vector suppression (Taning et al., 2020; Zhu and Palli, 2020; Zotti et al., 2018). Moreover, this technique allows identification of insect gene function with regard to interactions with phytoplasmas for transmission.

RNAi is a sequence-specific mechanism of eukaryotes, that regulates gene expression and provides a natural defence against nucleic acids of transposons or viruses (Taning et al., 2020). As a molecular tool, RNAi allows specific post-transcriptional silencing of target genes, and is a powerful tool for understanding gene function and regulation (Zotti et al., 2018). RNAi efficiency varies among insect orders (being Lepidoptera, Diptera, Hymenoptera, and Hemiptera the most studied ones), and also within species in the same order (Zhu and Palli, 2020). Many uncertainties remain on RNAi mechanisms occurring in insects, especially in non-model species, although RNAi has been reported for several hemipteran virus vectors (Chen et al., 2015; Kanakala and Ghanim, 2016; Matsumoto and Hattori, 2016; Xue et al., 2020) as well as for the phytoplasma vector *Euscelidius variegatus* Kirschbaum (Hemiptera: Cicadellidae (Abbà et al., 2019).

*Euscelidius variegatus* is a natural vector of chrysanthemum yellows (CYp) isolate of the *‘Candidatus* Phytoplasma asteris’, and an efficient vector of Flavescence dorée phytoplasma (FDp) under laboratory conditions (Galetto et al., 2018). *‘Candidatus* Phytoplasma asteris’ includes isolates associated with over 100 economically relevant plant diseases worldwide and is the most diverse and widespread phytoplasma group (Lee, 2004). The CYp isolate is associated with a disease of ornamental plants in Northwestern Italy (Conti et al., 1988). Flavescence dorée phytoplasma is a quarantine pest and causes an economically important grapevine disease representing one of the major threats to southern European viticulture (EFSA Panel on Plant Health PLH, 2014). Under field conditions, *E. variegatus, Macrosteles quadripunctulatus* Kirschbaum, and *Euscelis incisus* Kirschbaum are efficient vectors of CYp (Bosco et al., 2007; Conti et al., 1988), whereas FDp is transmitted to grapevine by *Scaphoideus titanus* Ball (EFSA Panel on Plant Health PLH, 2014). Compulsory insecticide treatments against the latter species as well as roguing of infected grapevines and their replacement with new grafted cuttings are the main preventive strategies to control the disease (EFSA Panel on Plant Health PLH et al., 2016). There is urgent demand for new and more sustainable management strategies to cope with economically important phytoplasma diseases such as FD. For this purpose, knowledge gaps on mechanisms exploited by phytoplasmas to colonize vectors must be filled. A laboratory model has been established to manage the FDp infection cycle using the herbaceous *Vicia faba* as host plant and the polivoltine leafhopper *E. variegatus* as laboratory vector (Caudwell et al., 1972), thus overcoming difficulties in dealing with a woody host (grapevine) and a monovoltine vector (*S. titanus).*

Phytoplasmas lack cell walls and therefore proteins exposed on the pathogen membrane are in direct contact with the host proteins. Several lines of evidence indicate that phytoplasma membrane proteins are crucial during the infection process of either plant or insect hosts, as reviewed in (Rossi et al., 2019). In particular, previous studies demonstrated that actin and ATP synthase α/β subunits of different insect vectors interact *in vitro* with the phytoplasma Antigenic membrane protein (Amp) of *‘Ca.* P. asteris’, which is required for pathogen transmission *in vivo* (Galetto et al., 2011; Rashidi et al., 2015; Suzuki et al., 2006). RNAi has been successfully applied to silence muscle actin and ATP synthase β genes of *E. variegatus,* following abdominal microinjection of dsRNAs (Abbà et al., 2019). The latter protein is part of the F1-F0 ATP synthase complex, a large multi-subunit mitochondrial enzyme that uses the proton gradient generated by the respiratory chain to synthesize ATP, whose general structure is highly conserved throughout evolution (Leyva et al., 2003).

The present work aims at characterizing gene silencing in *E. variegatus,* its systemic spread in the insect body, duration, small RNA (sRNA) profile and effects on the corresponding protein level. Moreover, we demonstrate that vector ATP synthase β silencing suppresses phytoplasma multiplication, a further proof of the functional role of this gene in the phytoplasma infection of the insect.

## 2. Materials and Methods

### 2.1. Insect rearing and phytoplasma isolates

*Euscelidius variegatus* were originally collected in Piedmont region of Italy and continuously reared on oat, *Avena sativa* (L.), inside plastic and nylon cages in growth chambers at 20–25 °C with a L16:D8 photoperiod (Rashidi et al., 2014). To obtain same age *E. variegatus* adults, about two weeks before the experiments, the required amount of 4^th^ and 5^th^ instar nymphs were taken from the main rearing and caged altogether on oats. Newly emerged adults were then injected with dsRNAs.

Chrysanthemum yellows phytoplasma (CYp, 16SrI-B), belonging to the *‘Candidatus* Phytoplasma asteris’ species, was isolated in the Italian Riviera (Liguria region) and maintained by insect transmission on daisy, *Chrysanthemum carinatum* Schousboe (Rashidi et al., 2014). Flavescence dorée phytoplasma (FDp, 16SrV-C) was identified in a vineyard of the Piedmont region of Italy, and transmitted to broad bean, *Vicia faba* L., by the natural vector *S. titanus* previously fed on infected grapevines; FDp was then maintained on broad bean by insect transmission, using the laboratory vector *E. variegatus* (Galetto et al., 2014). Daisies, broad beans, and oats were all grown from seed in greenhouses. For each acquisition access period (AAP), the sanitary status of source plants was confirmed by symptom observation and PCR diagnosis as previously described (Rashidi et al., 2014).

### 2.2. Synthesis and delivery of dsRNAs

The complete coding sequence of *E. variegatus* ATP synthase *β* (target mRNA) used in this work can be found in the TSA sequence database (BioProject: PRJNA393620) at NCBI under the accession number GFTU01013594.1 (Galetto et al., 2018).

The molecules of dsRNAs targeting *E. variegatus* ATP synthase *β* (dsATP) and the gene sequence of the green fluorescent protein (dsGFP), used as negative control, were synthetized as detailed in (Abbà et al., 2019). Briefly, plasmids harbouring target sequences under T7 promoter regulation were *in vitro* transcribed with MEGAscript RNAi Kit (Thermo Fisher Scientific), purified with ssDNA/RNA Clean and Concentrator (Zymo Research), eluted in Tris–EDTA buffer (10mM Tris–HCl, 0.1mM EDTA, pH 8.5) and quantified with a Nanodrop ND-1000 spectrophotometer (Thermo Fisher Scientific).

Newly-emerged adults were anaesthetized with CO2 and microinjected between two abdominal segments under a stereomicroscope using a fine glass needle connected to a Cell Tram Oil microinjector (Eppendorf). Insects were microinjected with 0.5 μL of dsRNAs at the concentration of 160 ng μL^-1^. Groups of injected insects were then caged on oat plants and monitored daily until the end of the experiments. Dead insects were removed periodically.

### 2.3. RNA extraction, cDNA synthesis, and gene expression analysis

Total RNAs were extracted from single *E. variegatus* adult (separated head and body or whole insect) at different times after dsRNA injection. The samples were frozen with liquid nitrogen, crushed with a micropestle in sterile Eppendorf tubes, and homogenized in 0.5 mL Tri-Reagent (Zymo Research). Samples were centrifuged 1 min at 12000 g at 4°C and RNAs were extracted from supernatants with Direct-zol RNA Mini Prep kit (Zymo Research), following manufacturer’s protocol and including the optional DNAse treatment step. Concentration, purity, and quality of extracted RNA samples were analysed in a Nanodrop ND-1000 spectrophotometer (Thermo Fisher Scientific).

Quantitative RT-PCR (qRT-PCR) was used to quantify the ability of the injected dsRNAs to knockdown target mRNA (ATP synthase *β*) in insects collected at 15, 22 and 37 days post injection (dpi), analysed as whole body (22 and 37 dpi) or as separated head and body (15 dpi). For each sample category, at least 10 biological replicates with balanced sex ratio were analysed to compare levels of ATP synthase *β* transcript between insects injected with dsATP and dsGFP. For each sample, cDNA was synthesized from total RNA (1 μg) with random hexamers using a High Capacity cDNA reverse transcription kit (Applied Biosystems). The resulting cDNA was used as a template for qPCR in a 10 μL volume mix, containing 1× iTaq Universal Sybr Green Supermix (Bio-Rad) and 300 nM of each primer. All primer pairs used in this work are listed in Supplementary Table S1. Samples were run in duplicate in a CFX Connect Real-Time PCR Detection System (Bio-Rad). Cycling conditions were: 95 °C for 3 min, and 40 cycles at 95 °C for 15 s and 60 °C for 30 s of annealing/extension step. The specificity of the PCR products was verified by melting curve analysis for all samples. No-template and no-reverse transcribed controls were always included in each plate. Primers targeting glutathione S-transferase and elongation factor-1α were used as internal controls to normalize the cDNA among samples. Normalized expression levels of each target gene for each sample were calculated by CFXMaestro™ Software (Bio-Rad). As no significant gender-based differences were observed, male and female samples were pooled together.

In order to check for the presence of the whole dsGFP sequence in dsGFP-injected insects at 22 dpi, cDNA was synthesized from total RNA (200 ng) using the High Capacity cDNA reverse transcription kit (Thermo Fisher Scientific Inc., MA, USA). Qualitative RT-PCR was conducted with the same primers used for the dsGFP synthesis without the T7 tail (Supplementary Table S1).

### 2.4. Library construction, sequencing, and bioinformatic analyses of small RNAs

Total RNAs extracted from whole insect bodies sampled on the 22^nd^ day after the injection of either dsATP or dsGFP were sent to Macrogen Inc. (South Korea) for small RNA (sRNA) library construction and sequencing. Three biological replicates were prepared for each condition, each containing a pool of six individuals: Eva_ATP1, Eva_ATP2 and Eva_ATP3 libraries for insects injected with dsATP and Eva_GFP1, Eva_GFP2 and Eva_GFP3 libraries for insects injected with dsGFP. Briefly, libraries were prepared with the Illumina TruSeq Small RNA Library construction kit (Illumina Inc., San Diego, CA, USA) and sequenced by the Illumina HiSeq2500 in rapid run mode 1×50bp. Raw reads were trimmed from adapters with Cutadapt, version 1.18 (Martin, 2011), and filtered out according to a) quality and length (minimum 16 nt; maximum 30 nt) with Reformat, version 37.71 (part of BBMap suite) (Bushnell, 2014) and b) mapping onto the *E. variegatus* ATP synthase *β* sequence. Bowtie version 1.1.2 software (Langmead, 2010) with no mismatch allowed in the alignment was used to establish sRNA abundance profiles of the six sequenced samples. Following alignment, the resulting SAM files were converted to BAM format, sorted by position and indexed using SAMtools version 1.9 (Li et al., 2009) and visualized with the Integrative Genomics Viewer (IGV) (Robinson et al., 2017). sRNA counts were normalized for differences in sequencing depths to account for the technical differences across samples.

### 2.5. Western blots

This procedure was used to quantify the expression of the ATP synthase *β* protein, in *E. variegatus* specimens collected at 4, 6, 8, 12 and 15 days post injection (dpi) of silencing dsRNAs. At each sampling date, and for each group of insects (dsATP vs. dsGFP), four to six insect samples were singly analyzed. Single insects were homogenized in 1.5 mL tube with micro-pestle in 100 μL of Rx Buffer (0.1% Triton X-100, 100 mM KCl, 3 mM NaCl, 3.5 mM MgCl2, 1.25 mM EGTA, and 10 mM Hepes, pH 7.3) (Suzuki et al., 2006), sonicated for 1 min at RT and centrifuged for 1 min at 13000 g. The supernatant was recovered and an aliquot was quantified in a UV-vis spectophotometer with Bradford reagent (Bio-Rad) along with a standard curve of known dilutions of Bovine Serum Albumin (BSA) dissolved in Rx Buffer. For each sample, 1 μg of total proteins was loaded onto 12% polyacrylamide gel, together with 3 μL of Sharpmass VI Prestained Protein Marker (EuroClone) and 5 μL of Unstained SDS-PAGE Standards, broad range (Bio-Rad). Gels were either stained with colloidal Coomassie stain (Candiano et al., 2004) or blotted on a polyvinylidene difluoride (PVDF) membrane. Membranes were blocked for 30 min with 3% BSA dissolved in Tris-buffered saline with 0.1% Tween (BSA-TBST) and incubated overnight at 4 °C with primary antibody (ab43177 chicken-developed anti-ATP synthase *β,* Abcam plc) diluted 1:5000 in BSA-TBST. Blots were then washed four times with BSA-TBST, incubated for 2 h with horseradish peroxidase (HRP)-conjugated secondary antibody (12-341 Rabbit anti-chicken RAC-HRP, Sigma-Aldrich) diluted 1:15000 in BSA-TBST, washed four times with TBST, and developed with West Pico SuperSignal chemiluminescent substrate (Pierce) in a VersaDoc 4000 MP system (Bio-Rad).

Quantity One 1-D Analysis Software (Bio-Rad) was used to estimate band intensities of each sample, expressed as intensity/mm^2^.

### 2.6. Phytoplasma acquisition, detection, and quantification

Following injection of dsRNAs (targeting either GFP as a control or ATP synthase β), insects were allowed to acquire phytoplasma by feeding on infected plants for few days (two and four days of acquisition access period, AAP, for CYp and FDp, respectively), then isolated on oat plants for a short latency (five and three days for CYp and FDp, respectively), and thus collected for molecular analyses at seven days post acquisition (dpa), as depicted in Figure 1.

**Figure 1.**
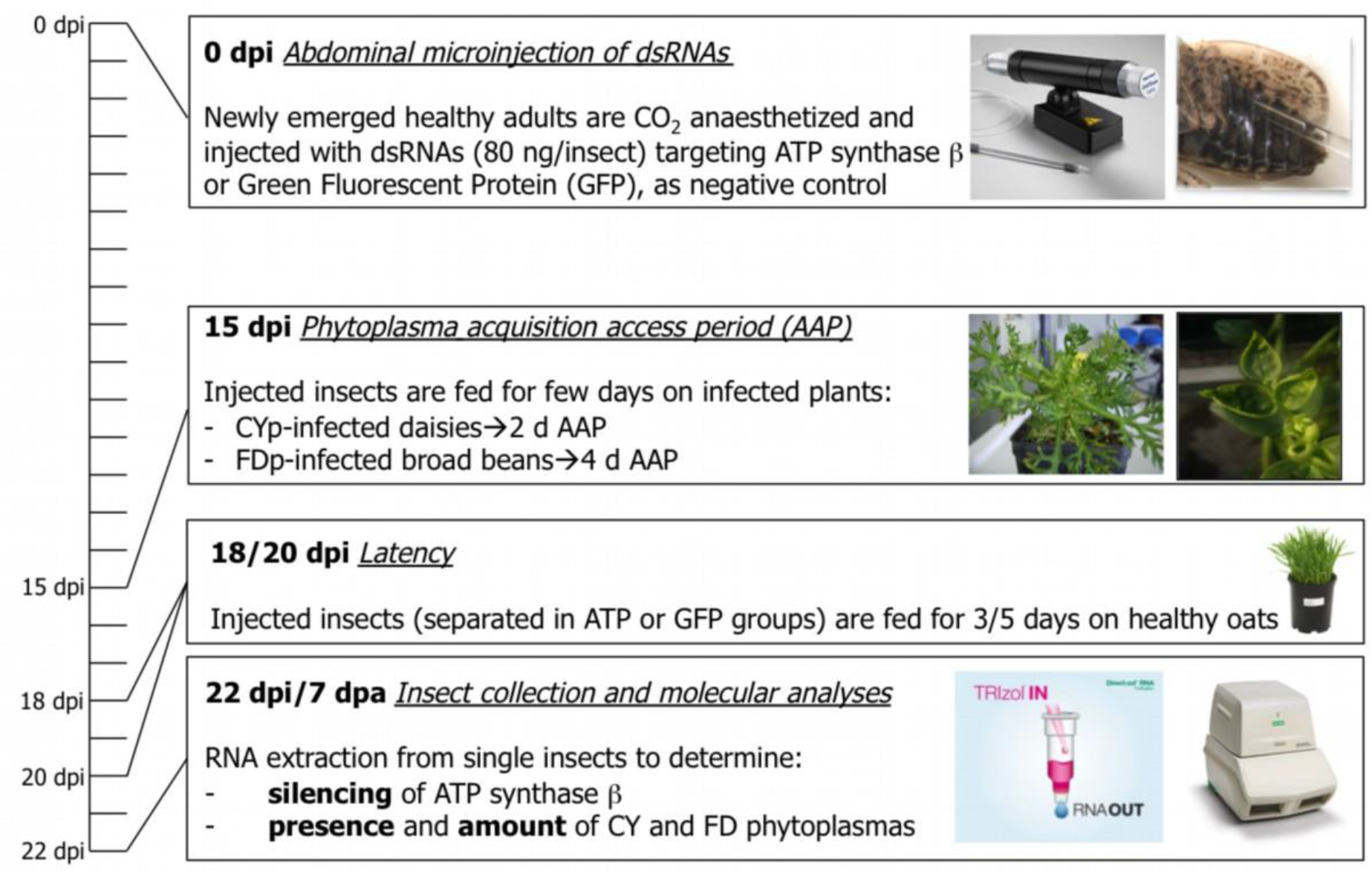
Methodology of phytoplasma acquisition experiments. dpi: days post injection; dpa: days post acquisition.

A multiplex quantitative RT-PCR (qRT-PCR) was used to detect phytoplasma presence and measure pathogen load in insects injected with dsATP or dsGFP. About 40 biological replicates were analysed for each dsRNA-injected group. Universal phytoplasma primers CYS2Fw/Rv and FAM-labelled TaqMan CYS2Probe designed on ribosomal 16SrRNA gene sequence were used to detect CYp or FDp (Marzachí and Bosco, 2005). Primers GapFw632/GapRv819 and HEX-labelled TaqMan GapEvProbe were used to measure *E. variegatus* glyceraldehyde-3-phosphate dehydrogenase (GAPDH) transcript, chosen as endogenous normalization control (Ottati et al., 2020). The sequences of primers and probes are detailed in Supplementary Table S1. For each sample, cDNA was synthesized as described above and one μL was used as template in a multiplex reaction mix of 10 μL total volume, containing 1x iTaq Universal Probes Supermix (Bio-Rad), 300 nM of each of the four primers and 200 nM of each of the two TaqMan probes. Each sample was run in triplicate in a CFX Connect Real-Time PCR Detection System (BioRad). Cycling conditions were 95°C for 3 min and 40 consecutive cycles at 95°C for 10 s as denaturing step followed by 30 s at 60°C as annealing/extension step. In each qPCR plate, four serial 100-fold dilutions of pGem-T Easy (Promega) plasmids, harbouring the target phytoplasma and insect genes, were included for relative quantification of the pathogen loads. For both plasmid standard curves, dilutions included in plates ranged from 10^8^ to 10^2^ target copy numbers per μL. Dilution series of both plasmids were used to calculate qPCR parameters (reaction efficiency and R^2^). Mean copy number of phytoplasma ribosomal transcripts were used to express pathogen amount as CYp or FDp 16SrRNA/insect GAPDH transcript. The experiment was repeated twice for each phytoplasma species.

### 2.7. Statistical analyses

SigmaPlot version 13 (Systat Software, Inc., www.systatsoftware.com.) was used for statistical analyses. Kruskal Wallis test, followed by Dunn’s method as pairwise multiple comparison procedure, was used to compare ATP synthase β transcript levels measured in heads or bodies of insects injected with dsGFP or dsATP. Mann-Whitney rank sum test was used to compare mean levels of ATP synthase β transcripts measured in insects injected with dsGFP or dsATP as well as to compare the mean phytoplasma loads quantified in the same insect groups. To compare pixel intensities in WB images between insects injected with dsGFP or dsATP t-test was used.

## 3. Results

### 3.1. Silencing of ATP synthase β is systemic and long-lasting

Abdominal microinjection of ATP synthase β dsRNAs into *E. variegatus* adults caused a significant strong decrease in the expression of the target gene, with a reduction ranging from 50 to over 130 times in comparison with the corresponding dsGFP-treated controls, according to the different experiments (Figure 2 and Supplementary Table S2). At 15 days post injection (dpi) silencing of the target was recorded both at the injection site (abdomen) and in the head, indicating systemic diffusion of the silencing signals. Indeed, significant reduction of ATP synthase β transcription was observed both in dissected heads and abdomens of insects injected with dsATP in comparison with those of insects injected with dsGFP (Figure 2a; Kruskal-Wallis, H=29.545, P<0.001, Dunn’s Method).

**Figure 2.**
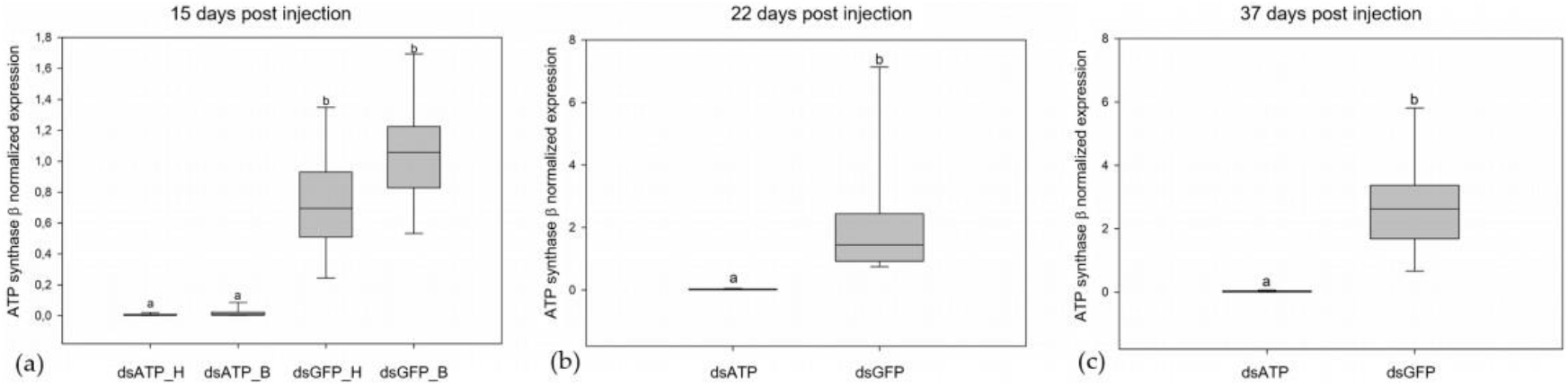
Transcript level of ATP synthase β in insects injected with dsRNAs. Transcript level of ATP synthase β in insects injected with dsRNAs targeting ATP synthase β (dsATP) or Green Fluorescent Protein (dsGFP) at 15 (**a**), 22 (**b**) and 37 (**c**) days post injection (dpi). At 15 dpi (**a**), **t**ranscript levels were measured separately in heads (_H) and abdomens (_B) of the different treated insect groups. At 22 **(b)** and 37 **(c)** dpi, **t**ranscript levels were measured in the whole body. The median is depicted as the line across the box, the box indicates the 25^th^ and 75^th^ percentiles, whiskers represent the 90^th^ and 10^th^ percentiles and different letters indicate significant differences between treatments.

Transcriptional silencing also showed long-lasting effects, as ATP synthase β transcript levels remained significantly lower in insects injected with dsATP than in those treated with dsGFP, at least up to 37 dpi (Figure 2b and c; Mann-Whitney, U=0.000, T=10428, P<0.001 at 22 dpi and U=0.000, T=105, P<0.001 at 37 dpi).

### 3.2. Generation of dsRNA-derived siRNAs

Total sRNA reads mapping to the dsATP sequence in Eva_ATP libraries (Figure 3a) and the dsGFP sequence in Eva_GFP libraries (Figure 3b) showed the same length distributions, the majority being in the 19-22 nt range with a peak at 21 nt (Supplementary Table S3). These observations confirmed that the silencing mechanism was still active at least until 22 dpi.

**Figure 3.**
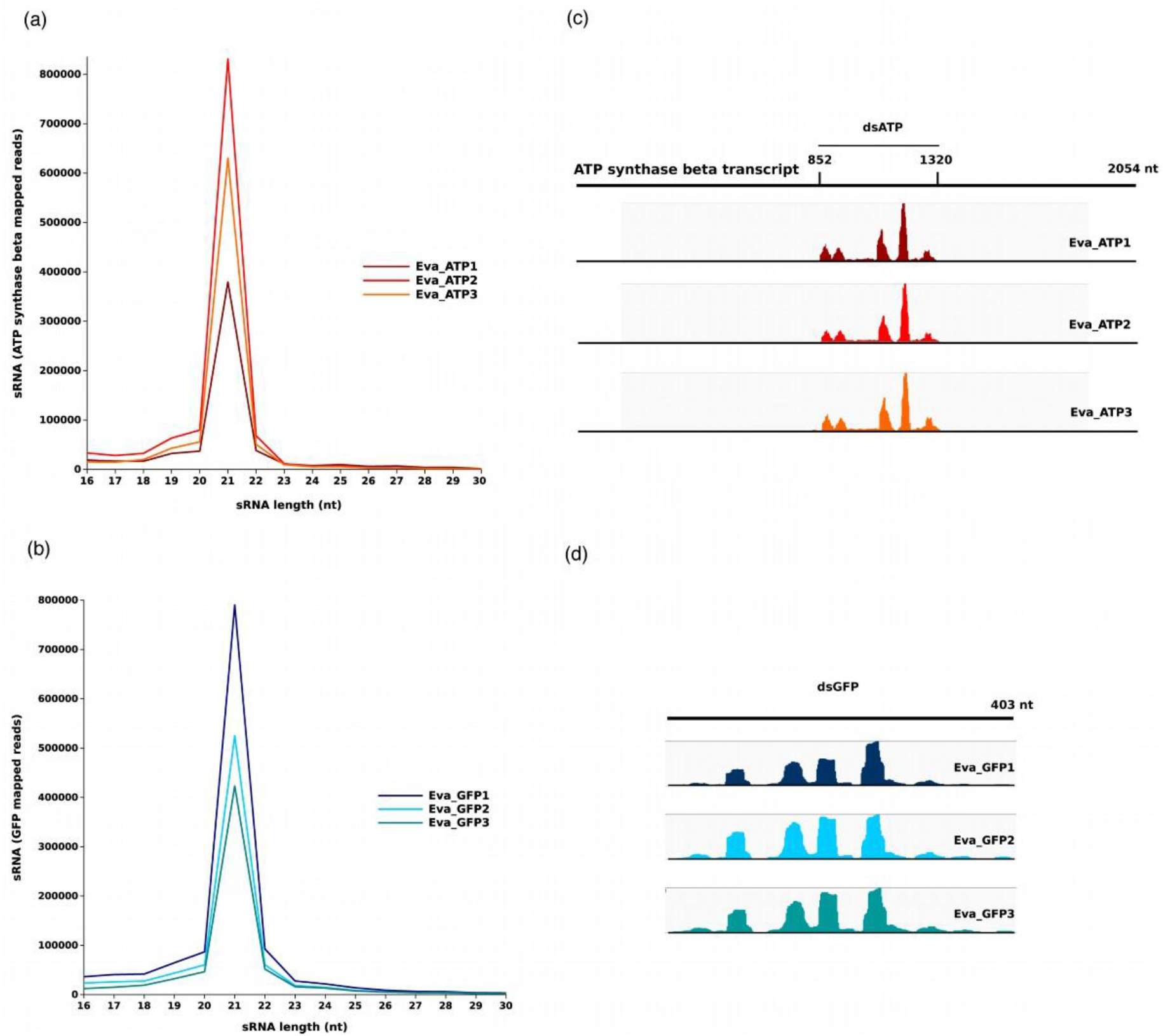
Analysis of smallRNAs profile. sRNA length distribution and mapping to ATP synthase β and Green Fluorescent Protein (GFP) in dsATP injected and dsGFP injected insects, respectively, at 22 days post injection (dpi). **(a)** Length distribution of sRNA mapping to the ATP synthase β transcript in the three libraries obtained from dsATP-injected insects. **(b)** Length distribution of sRNA mapping to the GFP in the three libraries obtained from dsGFP-injected insects **(c)** sRNA distribution over the whole length of the ATP synthase β transcript in the three libraries obtained from dsATP-injected insects. No sRNAs mapped outside the region corresponding to the injected dsATP. **(d)** sRNA distribution over the dsGFP sequence in the three libraries obtained from dsGFP-injected insects.

In Eva_ATP libraries, sRNA reads mapped exclusively to the fragment of the ATP synthase β transcript corresponding to the injected dsATP (Figure 3c). In both Eva_ATP and Eva_GFP libraries, sRNA covered the whole length of the corresponding dsRNA targets, with an uneven distribution. Two main hotspots could be observed between nt 247 and 362 in the dsATP sequence (Figure 3c), whereas in the dsGFP sequence, two peaks were clearly visible between nt 202 and 278 (Figure 3d). In both conditions hotspots were made up mostly of 21 nt-long sRNAs.

It is noteworthy that in four out of 12 dsGFP-injected insects collected at 22 dpi it was still possible to amplify the whole dsGFP sequence (data not shown), thus revealing unexpected long-term stability of dsRNAs. An analogous RT-PCR was not carried out on dsATP-injected insects, as in that case, it was impossible to discriminate the presence of intact dsATP from residual ATP synthase β transcripts.

### 3.3. Silencing of ATP synthase β gene affects the amount of encoded protein

Insects collected at 4, 6, 8, 12, and 15 dpi from the two treatments (dsATP vs. dsGFP) were analyzed by Western blot to quantify the amount of the corresponding ATP synthase *β* protein. The anti-ATP synthase β antibody recognized a protein of about 55 kDa (Supplementary Figure S1), as expected from *in silico* translation of the coding sequence (theoretical pI/Mw: 5.20/55.7 kDa). The protein expression level was similar in the two insect groups up to 12 dpi, whereas at 15 dpi the protein amount detected in the six insects injected with dsATP was about 2.5 times lower (t-test, t=4.510, P<0.001) than that observed in the six insects injected with dsGFP (Figure 4a, b, and Supplementary Table S4). Coomassie staining of the same protein samples separated by SDS-PAGE confirmed that equal amounts of proteins from the two insect groups were loaded into gels (Supplementary Figure S1).

**Figure 4.**
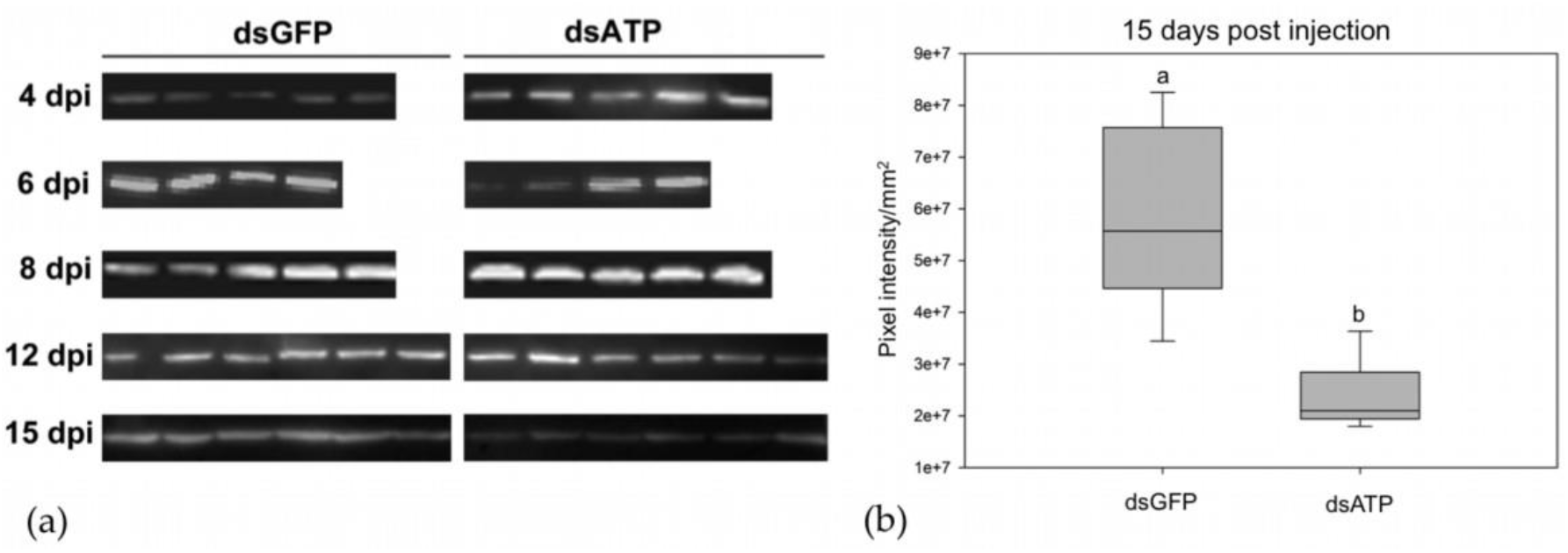
Protein level of ATP synthase β in insects injected with dsRNAs. **(a)** Protein expression of ATP synthase β analysed by Western blots on *Euscelidius variegatus* insects (one insect sample per lane) injected with dsRNAs targeting ATP synthase β (dsATP) or Green Fluorescent Protein (dsGFP) at 4, 6, 8, 12 and 15 days post injection (dpi). **(b)** Pixel intensity/mm^2^ of Western blots bands depicted in **(a)** measured in insects analysed at 15 dpi. This parameter is the sum of the intensity of each pixel calculated by Quantity One 1-D Analysis Software (BioRad) included in the band boundary manually defined. The median is depicted as the line across the box, the box indicates the 25^th^ and 75^th^ percentiles, whiskers represent the 90^th^ and 10^th^ percentiles and different letters indicate significant differences between treatments.

### 3.3. Silencing of ATP synthase β reduces pathogen multiplication, while it has no effect on prevalence of infected insects

Insects injected with either dsATP or dsGFP successfully acquired either CYp or FDp during feeding on infected plants, as all analysed insects tested positive in RT-qPCR for the presence of both pathogens (Table 1).

**Table 1.** Phytoplasma acquisition by insects after silencing of ATP synthase β. Acquisition of phytoplasmas by *Euscelidius variegatus* insects, sampled at 22 days post injection of dsRNAs targeting ATP synthase β or green fluorescent protein (GFP) and at 7 days post acquisition of chrysanthemum yellows (CYp) or Flavescence dorée (FDp) phytoplasmas, as detailed in methodology depicted in Figure 1.

| Target of dsRNAs | Phytoplasma acquired post dsRNA injection | Phytoplasma positive/analysed insects |
| --- | --- | --- |
| ATP synthase $\beta$ | CYp | 42/42 |
| GFP | CYp | 43/43 |
| ATP synthase $\beta$ | FDp | 39/39 |
| GFP | FDp | 36/36 |

On the contrary, significant differences were observed in phytoplasma mean quantities measured in insects injected with dsATP compared with those measured in insects injected with dsGFP, for both CYp and FDp (Figure 5 and Supplementary Table S5; Mann-Whitney U=597, T=1500, P=0.007 for CYp and U=466, T=1604, P=0.013 for FDp). Mean CYp and FDp amounts in dsATP insects were five and four time lower than those measured in corresponding dsGFP-injected specimens, indicating that silencing of ATP synthase β reduced multiplication rates of both pathogens. The copy number of CYp 16S RNA per insect GAPDH transcript ranged from 1.83E-06 to 7.56E-02 with a median of 1.79E-03 in dsATP-injected insects and from 1.08E-05 to 2.50E-01 with a median of 4.88E-03 in dsGFP-injected ones. The copy number of FDp 16S RNA per insect GAPDH transcript ranged from 3.67E-07 to 3.34E-01 with a median of 1.89E-03 in dsATP insects and from 8.50E-06 to 5.45E-01 with a median of 1.24E-02 in dsGFP insects.

**Figure 5.**
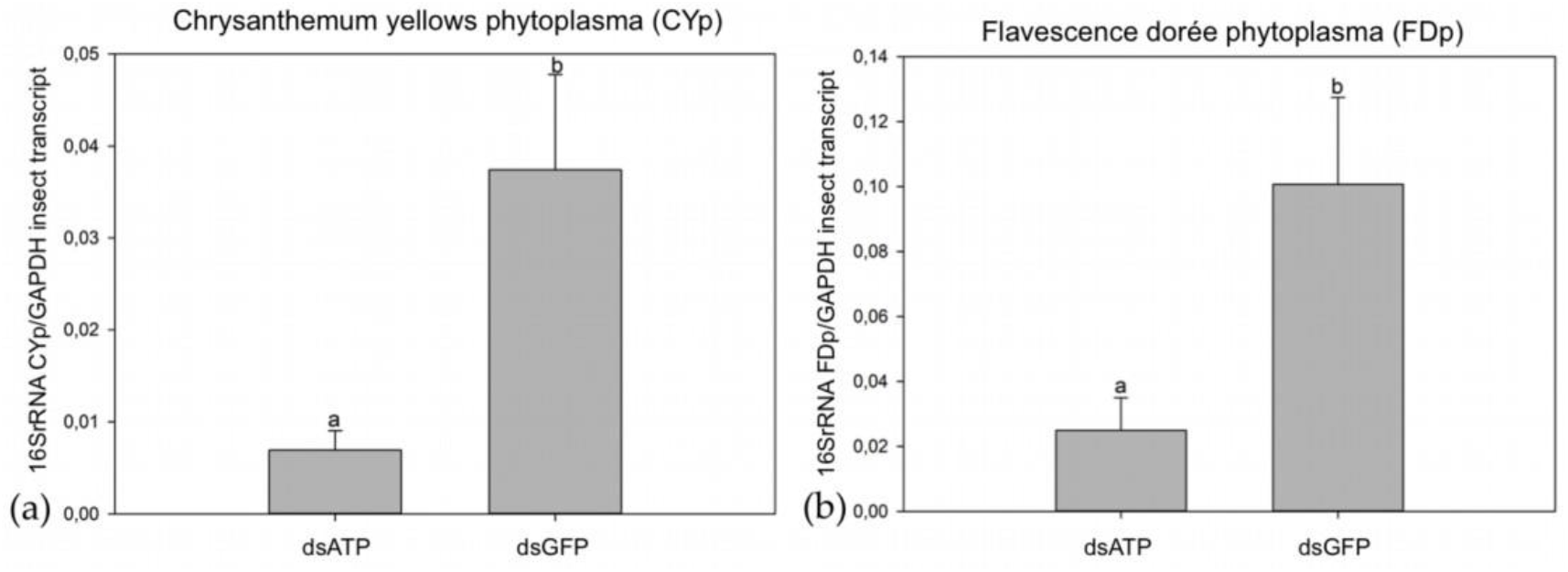
Phytoplasma amount in insects after silencing of ATP synthase β. Mean phytoplasma 16S RNA/insect GAPDH transcript ± standard error of the mean (SEM) measured in *Euscelidius variegatus* insects, sampled at 22 days post injection of dsRNAs targeting ATP synthase β (dsATP) or green fluorescent protein (dsGFP). Twenty-two days post injection correspond to 7 days post acquisition of chrysanthemum yellows (CYp, panel **a**) or Flavescence dorée (FDp, panel **b**) phytoplasmas. Different letters indicate significant differences in mean phytoplasma amount.

## 4. Discussion

The present work shows that silencing of an insect vector gene, involved in interaction with the antigenic membrane protein of the transmitted phytoplasma, suppresses phytoplasma multiplication in the insect body. Since its discovery, RNAi has emerged as a powerful molecular tool for studying gene function, regulation, and interaction at the cell and organism levels. RNAi is particularly efficient in some insects, especially those included in the Coleoptera order, whereas species within Lepidoptera and Hemiptera may show variable attitudes in response to RNAi (Zhu and Palli, 2020; Zotti et al., 2018). Several efforts have been made using RNAi in hemipterans. In most of these cases, target genes for silencing were selected to achieve insect mortality, but in few instances RNAi was applied to interfere with vector ability. To this purpose, silencing of viral genes involved in interaction with vector proteins has been exploited, as revised in (Kanakala and Ghanim, 2016). In *Laodelphax striatellus,* silencing of *Argonaute* 2 enhanced the accumulation of the insect-specific Himetobi P picorna-like virus (Xu et al., 2014), and silencing of the cuticular protein CPR1 decreased transmission ability of rice stripe tenuivirus by the same planthopper (Liu et al., 2015). Silencing of *Bemisia tabaci* heat shock proteins disrupted interactions with tomato yellow leaf curl begomoviruses (Gotz et al., 2012; Ohnesorge and Bejarano, 2009). Here, silencing of ATP synthase β gene of the phytoplasma vector *E. variegatus*, besides confirming the vital role of this gene for efficient phytoplasma colonization of the vector body, represents the first attempt to disrupt phytoplasma-vector interactions, and paves the way to new strategies to interfere with phytoplasma transmission. Moreover, this mechanism seems to be conserved in two genetically unrelated phytoplasma species.

Gene silencing mediated by RNAi spread systemically in the leafhopper, as demonstrated by the reduction of ATP synthase β transcripts in the heads of insects after abdominal microinjection with the triggering dsRNA molecules. Two mechanisms of cellular uptake of dsRNAs have been identified in insects: Sid-like transmembrane channels, and clathrin-dependent endocytosis (Christiaens et al., 2020). SID-1 is a channel protein responsible for binding long dsRNAs for uptake by cells and is required for the systemic RNAi response in the nematode *Caenorhabditis elegans*, the organism in which RNAi has been first described (Fire et al., 1998). Sid-like genes have been identified in Coleoptera, Hemiptera, and Lepidoptera, but their role in cellular uptake has not been directly evidenced to date (Christiaens et al., 2020; Vélez and Fishilevich, 2018). Despite initially the *Sid-1* gene was not included among the key components of the *E. variegatus* RNAi machinery (Abbà et al., 2019), a manual analysis of the transcriptome confirmed its presence in this species. The Sid-1 coding sequence was, in fact, identified in the TSA entry GFTU01001708.1 from nt 7024 to nt 9375 on the reverse strand (Simona Abbà, personal communication). The presence of this gene in *E. variegatus* suggests its possible involvement in dsRNA uptake and in the spread of gene-silencing signals to adjacent tissues.

Injection of dsATP in *E. variegatus* determined a long-lasting reduction of the corresponding transcript by an average 100-fold compared to dsGFP controls, and the silencing was still efficient at least up to 37 days after the injection. Duration of RNAi is highly variable among insects. For example, the flour beetle *Tenebrio molitor* shows a robust RNAi response with gene suppression in virtually any cell type and also inheritance to progeny, whereas in *Drosophila* gene silencing is often short-termed and limited to few cell types (Bucher et al., 2002; Miller et al., 2008). Huge variability has been also described within Hemiptera order. In the pea aphid *Acyrthosiphon pisum* RNAi silencing of two marker genes starts one day after dsRNA injection, reaches its maximum at 5 days and stops at 7 days (Jaubert-Possamai et al., 2007). On the other hand, plant-mediated RNAi in the green peach aphid *Myzus persicae* has been shown to last up to 16 days and to be retained in the progeny when insects are continuously fed on dsRNA-producing transgenic *Arabidopsis thaliana*, whereas the silencing effect disappears within 6 days when the aphids are removed from the transgenic plants (Coleman et al., 2015). Parental RNAi has been demonstrated to occur also in the green rice leafhopper *Nephotettix cincticeps,* as injection of dsRNAs silences target genes at least up to 14 dpi in parent females as well as in the 1^st^ instar offspring nymphs (Matsumoto and Hattori, 2016). Parental RNAi has been confirmed in the triatomine bug *Rhodnius prolixus,* and interestingly the knockdown effects of three target genes persisted in this species up to 7 months after the dsRNA injection (Paim et al., 2013). Whether the long-term effect is still mediated by the primary dsRNA molecules initially injected or due to a secondary propagation of RNAi signal is unclear. Nevertheless, some hints may be deduced by the analysis of the sRNA pattern profiles obtained from silenced insects in comparison with GFP treated controls. Total sRNA reads mapping to the dsATP sequence in Eva_ATP libraries and the dsGFP sequence in Eva_GFP libraries showed length distributions of sRNA consistent with the processing of dsRNA into siRNA by the RNAi pathway, as previously observed in other insects (Santos et al., 2019). Interestingly, sRNA reads mapped exclusively to the fragment of the ATP synthase β transcript covering the injected dsATP. The lack of read coverage over the rest of the 2054 nt-long transcript pointed against the existence in *E. variegatus* of mechanisms of secondary siRNA synthesis similar to those observed in plants and nematodes (Christiaens et al., 2020). Consistently, the detection of intact dsGFP in some specimens at 22 dpi suggests that the long-term silencing could be due to an inefficient degradation of the injected dsRNAs, which continued to trigger the silencing machinery. Even though systemic RNAi is observed in insects, the specific mechanisms, genes involved in the spread of dsRNAs, and the kind of signal molecules (either dsRNA or siRNA) are yet to be unraveled (Vélez and Fishilevich, 2018). In the case of viral infection of *Drosophila,* transport of dsRNA through the insect body may occur via derived complementary viral DNAs used as template for *de novo* synthesis of secondary viral siRNAs in hemocytes and released in exosomes (Tassetto et al., 2017). Nanotube-like structures observed in *D. melanogaster* culture cells may also be one of the mechanisms of antiviral RNAi machinery transport between infected and non-infected cells (Karlikow et al., 2016).

Beside the reduction of the specific transcript, a significant decrease of the corresponding protein occurred starting from 15 days after dsRNA injection. The delay of the observed effect at protein level is consistent with the assembly of the mitochondrial respiratory complexes, which exploit oversynthesis of the nucleus-encoded proteins to provide an adequate molecule supply for the assembly process (Bogenhagen and Haley, 2020). Determining the effect of dsRNA injection on protein expression allowed us to optimize experiments with a proper time scale, as phytoplasma acquisition was performed only when the effect of gene silencing was evident at the protein level. Even though no effect was observed on phytoplasma acquisition efficiency, as all insects tested positive for the phytoplasma presence irrespective of dsATP or dsGFP injections, a significant lower pathogen amount was measured in silenced insects compared with the controls. ATP synthase β is already known to interact *in vitro* with the phytoplasma major Antigenic membrane protein (Galetto et al., 2011), function as a viral receptor (G.-F. Liang et al., 2015; Liang et al., 2010) and mediate the entry of viral particles into host arthropod cells (Fongsaran et al., 2014). Our data, are in line with this role of ATP synthase β. Interestingly, for a long time F1-F0 ATP synthase complex was thought to be exclusively located in the inner membrane of mitochondria, but more and more clues are pointing at the existence of ATP synthases on the outer face of plasma membranes of many cell lines and tissues of mammals, as reviewed in (Taurino and Gnoni, 2018), and arthropods (Fongsaran et al., 2014; Y. Liang et al., 2015; Lin et al., 2009). Moreover, the synthesis of extracellular ATP produced by F1-F0 complex expressed on plasma membranes has been documented for a wide variety of cells, mainly mammalian adipocytes, keratinocytes, endothelial, and hepatic cells (Taurino and Gnoni, 2018), but also primary cell lines of the yellow fever mosquito *Aedes aegypti* (Fongsaran et al., 2014) and hemocytes of the Pacific white shrimp *Litopenaeus vannamei* (Y. Liang et al., 2015). The ectopic expression of ATP synthase β in plasma membranes has been shown in midguts and salivary glands of *E. variegatus* (Galetto et al., 2011), so we may assume that the synthesis of extracellular ATP also occurs in *E. variegatus.* In addition, it is noteworthy that phytoplasmas lack the ATP synthetic pathway and strongly depend on their hosts for energy metabolism (Oshima et al., 2004). Therefore, a decrease in extracellular ATP caused by the silencing of ATP synthase β may impair the survival of phytoplasma cells and explain the observed reduction in the pathogen multiplication rate. The uptake of host-synthesized ATP seems to be a mechanism commonly shared by different phytoplasmas, as the phylogenetically distant CYp and FDp behaved similarly. Host-synthesized ATP plays a role during binding and successive infection process of the Pacific white shrimp by the white spot syndrome virus (Y. Liang et al., 2015). Phytoplasma transmission depends on the level of phytoplasma multiplication in the vector, as shown for CYp (Bosco et al., 2007; Galetto et al., 2009), and apple proliferation phytoplasma (Cainelli et al., 2007; Mayer et al., 2009), therefore the specific silencing of vector ATP synthase β may become part of a strategy aimed at reducing insecticide use against less efficient vectors. Indeed, the strong, long-lasting, and systemic nature of the observed gene silencing after a single dose of dsATP may theoretically identify RNAi as a suitable control strategy against phytoplasma insect vectors. From a practical point of view, dsRNA delivery to sap-sucking insects remains a big unsolved issue, despite new technologies and innovative approaches are envisaged (Christiaens et al., 2020). In the case, for example, of the Asian citrus psyllid *Diaphorina citri*, vector of citrus Huanglongbing, dsRNAs were successfully delivered by soaking insects in dsRNA-containing solutions (Killiny et al., 2014; Yu et al., 2017) and by feeding them on an optimized plant system (iPS) bioassay (Andrade and Hunter, 2017).

## 5. Conclusions

Silencing of ATP synthase β of *E. variegatus* decreased multiplication of two unrelated phytoplasmas in the vector body. As silencing of this gene resulted in the decrease of the corresponding protein, a role of ATP synthase β in mediating phytoplasma multiplication *in vivo* is suggested. This comprehensive study analyzed changes in transcript levels, protein expression, and sRNA profiles triggered by dsRNA injection. Gene silencing spreads systemically with unexpected long-lasting effects, which probably depend on insufficient dsRNA degradation rather than the production of secondary siRNAs, as suggested by the sRNA profiles. The presence of such a robust RNAi machinery is of immense interest, especially in the case of insect vectors of phytoplasmas, as it may be exploited as a possible tool to disrupt transmission and integrated in pest management programs to face phytoplasma diseases (Sattar and Thompson, 2016).

## Supplementary Materials

Table S1: Primers and probes, Table S2: Mean expression values of ATP synthase β transcript, Table S3: sRNA length distributions on dsRNA targets in the six sRNA libraries, Table S4: Pixel intensity values of Western blots images, Table S5: Phytoplasma detection and mean pathogen quantification values, Figure S1: Full length images of Western blots and corresponding SDS-polyacrylamide gels.

## Author Contributions

Conceptualization, L.G., S.A., D.B. and C.M.; methodology and investigation, L.G., S.A., M.Ro., M.Ri., and S.P.; writing —original draft preparation, L.G. and S.A.; writing – review and editing, M.Ro., S.P., D.B. and C.M.; funding acquisition, L.G., S.A., D.B. and C.M. All authors have read and agreed to the published version of the manuscript.

## Acknowledgments

This project has received funding from the European Union’s Horizon 2020 research and innovation programme under grant agreement No 773567. This research was also supported by Fondazione Cassa di Risparmio di Torino, Projects Siglofit (RF = 2016-0577) and FOotSTEP (RF = 2018-0678).

The authors thank Elena Zocca for providing plants for insect rearing, Flavio Veratti and Francesca Canuto for maintenance of insect colonies, Giulia Molinatto for helping in microinjection procedure.

---

**Supplementary Table S1.**
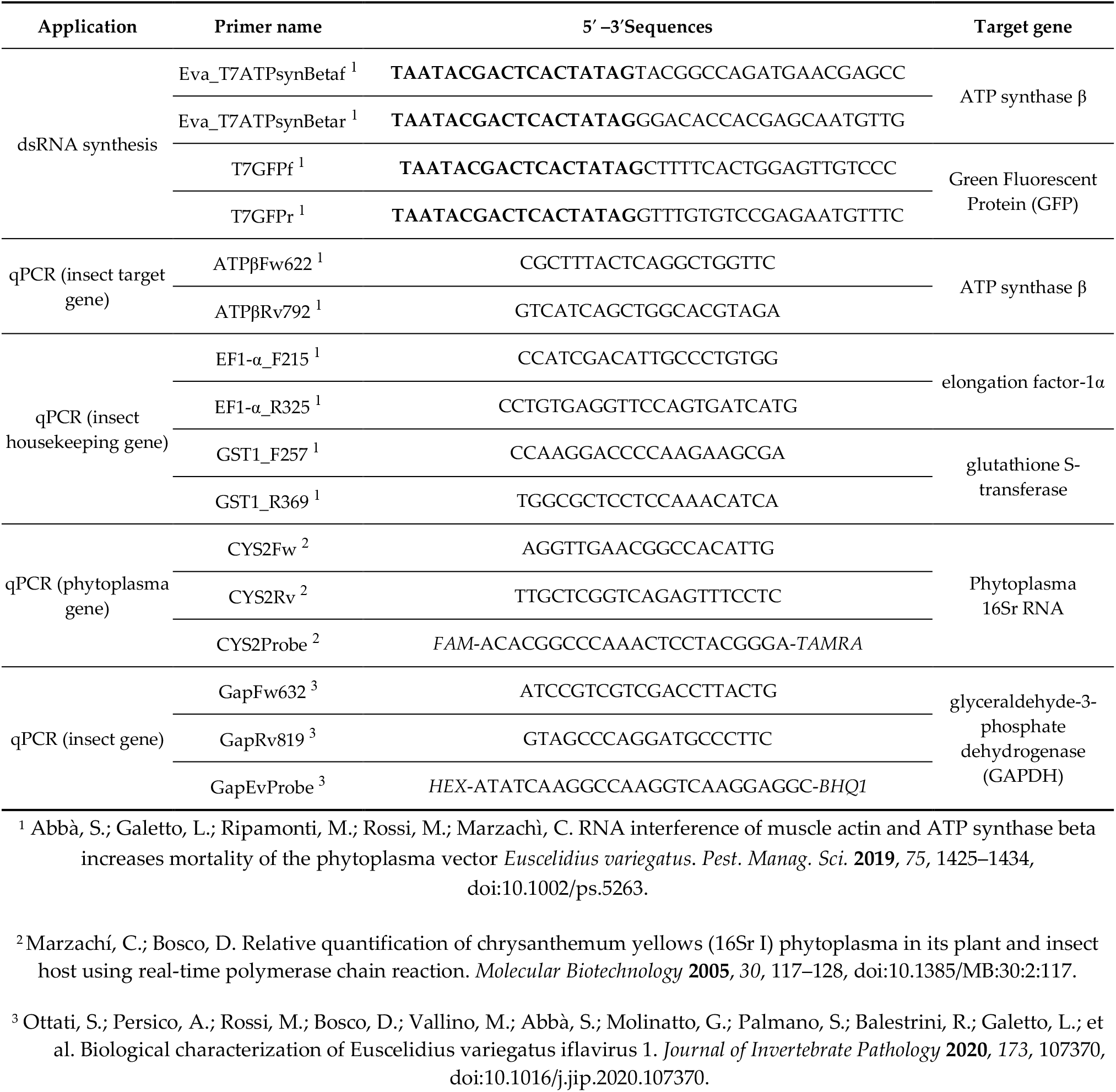
List of primers and probes used in this work. T7 promoter sequence is in bold. Fluorophore reporter and quencher are in italic.

**Supplementary Table S2.**
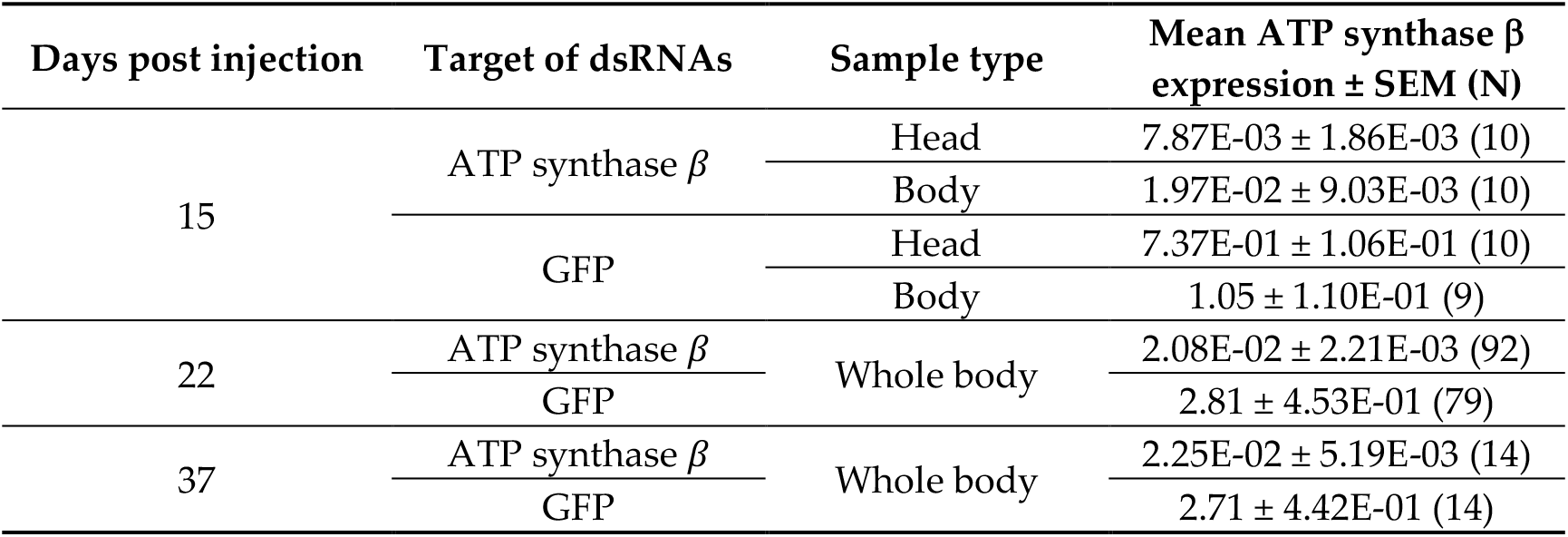
Mean normalized relative expression of ATP synthase β transcript ± standard error of the mean (SEM) measured in head/body/whole insect of *Euscelidius variegatus* samples at 15, 22 and 37 days post injection (dpi) of dsRNAs targeting ATP synthase β or green fluorescent protein (GFP).

| Days post injection | Target of dsRNAs | Sample type | Mean ATP synthase $\beta$ expression ± SEM (N) |
| --- | --- | --- | --- |
| 15 | ATP synthase $\beta$ | Head | 7.87E-03 ± 1.86E-03 (10) |
| | | Body | 1.97E-02 ± 9.03E-03 (10) |
| | GFP | Head | 7.37E-01 ± 1.06E-01 (10) |
| | | Body | 1.05 ± 1.10E-01 (9) |
| 22 | ATP synthase $\beta$ | Whole body | 2.08E-02 ± 2.21E-03 (92) |
| | GFP | | 2.81 ± 4.53E-01 (79) |
| 37 | ATP synthase $\beta$ | Whole body | 2.25E-02 ± 5.19E-03 (14) |
| | GFP | | 2.71 ± 4.42E-01 (14) |

**Supplementary Table S3.** Number of reads of different length (16-30 nt) mapping to dsRNAs targeting ATP synthase β (dsATP) or green fluorescent protein (dsGFP) in the six sRNA libraries analysed from the two groups of insects injected with either dsATP (Eva_ATP1, Eva_ATP2, Eva_ATP3) or dsGFP (Eva_GFP1, Eva_GFP2, Eva_GFP3).

| Reads mapping | Reads length | Library names |  |  |  |  |  |
| --- | --- | --- | --- | --- | --- | --- | --- |
|  |  | Eva_ATP1 | Eva_ATP2 | Eva_ATP3 | Eva_GFP1 | Eva_GFP2 | Eva_GFP3 |
| Mapping to dsATP | 16 | 21071 | 33198 | 16810 | 31 | 430 | 140 |
|  | 17 | 18519 | 28025 | 16425 | 41 | 384 | 146 |
|  | 18 | 18305 | 32338 | 22263 | 33 | 352 | 150 |
|  | 19 | 35983 | 63339 | 48217 | 26 | 621 | 291 |
|  | 20 | 41256 | 79437 | 62671 | 22 | 280 | 149 |
|  | 21 | 423242 | 830849 | 710122 | 68 | 420 | 339 |
|  | 22 | 42734 | 68026 | 57132 | 39 | 340 | 178 |
|  | 23 | 12293 | 11520 | 10636 | 13 | 223 | 145 |
|  | 24 | 8385 | 5630 | 5925 | 15 | 197 | 163 |
|  | 25 | 10718 | 6720 | 6084 | 24 | 300 | 241 |
|  | 26 | 6732 | 3370 | 3367 | 15 | 185 | 155 |
|  | 27 | 7646 | 3169 | 3419 | 22 | 320 | 208 |
|  | 28 | 4269 | 1955 | 1867 | 6 | 82 | 67 |
|  | 29 | 4477 | 1517 | 1560 | 6 | 106 | 68 |
|  | 30 | 1532 | 457 | 618 | 1 | 53 | 39 |
| Total dsATP mapped reads |  | 657162 | 1169550 | 967116 | 362 | 4293 | 2479 |
| Mapping to dsGFP | 16 | 7 | 61 | 5 | 36400 | 23395 | 15696 |
|  | 17 | 6 | 52 | 16 | 40724 | 26001 | 19222 |
|  | 18 | 3 | 67 | 13 | 41780 | 27448 | 24382 |
|  | 19 | 3 | 69 | 16 | 64683 | 43726 | 41178 |
|  | 20 | 9 | 39 | 20 | 86864 | 60390 | 59237 |
|  | 21 | 61 | 70 | 65 | 790116 | 525007 | 538637 |
|  | 22 | 12 | 32 | 13 | 92659 | 60695 | 66046 |
|  | 23 | 8 | 37 | 10 | 27650 | 18516 | 20514 |
|  | 24 | 4 | 28 | 8 | 21753 | 14643 | 16880 |
|  | 25 | 1 | 22 | 3 | 13859 | 8503 | 9623 |
|  | 26 | 1 | 13 | 2 | 8915 | 5416 | 6723 |
|  | 27 | 1 | 6 | 6 | 6502 | 4067 | 5236 |
|  | 28 | 2 | 11 | 2 | 6002 | 4254 | 4871 |
|  | 29 | 2 | 4 | 0 | 3829 | 2497 | 3008 |
|  | 30 | 1 | 3 | 3 | 3309 | 2290 | 2833 |
| Total dsGFP mapped reads |  | 121 | 514 | 182 | 1245045 | 826848 | 834086 |
| Total reads in library |  | 22281562 | 19966749 | 22495258 | 27404377 | 22204066 | 28271348 |

**Supplementary Table S4.** Mean intensity/mm^2^ of Western blots (WB) bands depicted in Figure 4 ± standard error of the mean (SEM) measured in *Euscelidius variegatus* insects, sampled at 15 days post injection of dsRNAs targeting ATP synthase β or green fluorescent protein (GFP). This parameter is the sum of the intensity of each pixel calculated by Quantity One 1-D Analisys Software (Bio-Rad) included in the band boundary manually defined.

| Days post injection | Target of dsRNAs | Mean intensity/mm<sup>2</sup> of WB bands ± SEM (N) |
| --- | --- | --- |
| 15 | ATP synthase $\beta$ | 2.37E07 ± 2.75E06 (6) |
|  | GFP | 5.83E07 ± 7.17E06 (6) |

**Supplementary Table S5.** Mean phytoplasma 16S RNAs/insect GAPDH transcript ± standard error of the mean (SEM) measured in *Euscelidius variegatus* insects, sampled at 22 days post injection of dsRNAs targeting ATP synthase β or green fluorescent protein (GFP) and at 7 days post acquisition of chrysanthemum yellows (CYp) or Flavescence dorée (FDp) phytoplasmas, as detailed in methodology depicted in Figure 1.

| Target of dsRNAs | Phytoplasma acquired post dsRNA injection | Mean phytoplasma quantity ± SEM (N) |
| --- | --- | --- |
| ATP synthase $\beta$ | CYp | 6.94E-03 ± 2.12E-03 (42) |
| GFP |  | 3.74E-02 ± 1.03E-02 (43) |
| ATP synthase $\beta$ | FDp | 2.49E-02 ± 1.00E-02 (39) |
| GFP |  | 1.01E-01 ± 2.68E-02 (36) |

**Supplementary Figure S1.**
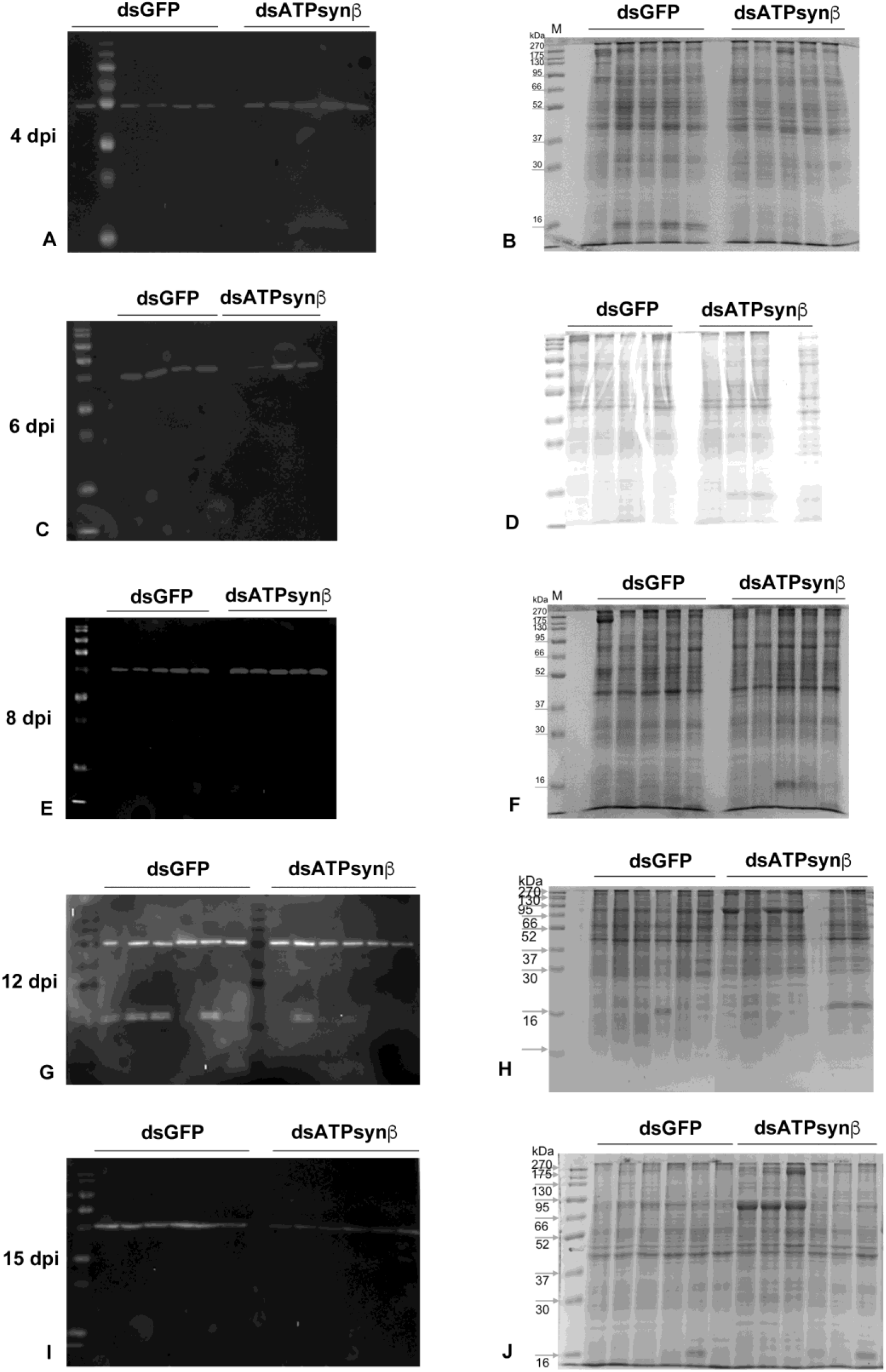
Full lenght images of Western blots (WB) (left column, A, C, E, G, I) depicted in Figure 4, developed with anti-ATP synthase β antibody, and corresponding Coomassie stained SDS-polyacrylamide gels (right column, B, D, F, H, J), run in parallel with WB on the same *Euscelidius variegatus* insect samples (one insect sample per single lane), analysed at 4, 6, 8, 12, 15 days post injection (dpi) of dsRNAs targeting ATP synthase β (dsATPsynβ) or green fluorescent protein (dsGFP).

## Notes

### Competing Interest Statement

The authors have declared no competing interest.

